# Spatial distribution and functional relevance of FGFR1 and FGFR2 expression for glioblastoma tumor invasion

**DOI:** 10.1101/2023.01.10.523434

**Authors:** Nawal Alshahrany, Ana Jimenez-Pascual, Florian A Siebzehnrubl

## Abstract

Glioblastoma is the most lethal brain cancer in adults. These incurable tumors are characterized by profound heterogeneity, therapy resistance, and diffuse infiltration. These traits have been linked to cancer stem cells, which are important for glioblastoma tumor progression and recurrence. The fibroblast growth factor receptor 1 (FGFR1) signaling pathway is a known regulator of therapy resistance and cancer stemness in glioblastoma. FGFR1 expression shows intertumoral heterogeneity and higher FGFR1 expression is associated with a significantly poorer survival in glioblastoma patients. The role of FGFR1 in tumor invasion has been studied in many cancers, but whether and how FGFR1 mediates glioblastoma invasion remains to be determined. Here, we investigated the distribution and functional relevance of FGFR1 and FGFR2 in human glioblastoma xenograft models. We found FGFR1, but not FGFR2, expressed in invasive glioblastoma cells. Loss of FGFR1, but not FGFR2, significantly reduced cell migration *in vitro* and tumor invasion in human glioblastoma xenografts. Comparative analysis of RNA-sequencing data of FGFR1 and FGFR2 knockdown glioblastoma cells revealed a FGFR1-specific gene regulatory network associated with tumor invasion. Our study reveals new gene candidates linked to FGFR1-mediated glioblastoma invasion.

## Introduction

Glioblastomas (GBMs) are the most common and malignant type of brain tumor in adults ^1^. Histopathological hallmarks of GBM include highly infiltrating cancer cells with increased mitotic activity, endothelial proliferation, and necrosis ^2, 3^. GBM accounts for 50% of gliomas with most incidences occurring between the ages of 40-60, and it is higher in men than women ^4,5^.

Current first-line treatment regimens include surgical resection followed by radiotherapy and/ or chemotherapy (usually temozolomide (TMZ)) ^6-8^. In recent years, molecular targeted therapies including tyrosine kinase inhibitors and immunotherapy showed promising results in treating GBM patients. Despite the aggressive treatment, patients with GBM usually have low median survival of 7–15 months from the time of diagnosis. This is because most patients develop recurrent disease 6–9 months after treatment making clinical management challenging ^9, 10^. It is not clear what causes GBM recurrence, however, it has been suggested that it could be due to the infiltrative nature of GBM cells, intratumoral heterogeneity, and the presence of a glioblastoma cancer stem cell (GSC) population ^11, 12^.

GSCs are a small population of tumor cells that can initiate and drive tumor development. GSCs have been shown to be resistant to both chemo and radiotherapies and more invasive than other cells of the same tumor ^13, 14^. GSCs are further able to thrive in harsh microenvironments and being unaffected by strict cellular checkpoints on their self-renewal and differentiation ^15, 16^. Several signaling pathways which promote their stemness and growth similar to neural stem cells (NSC) have been identified in GSCs including Sonic hedgehog, Notch, and Wnt ^16, 17^. Furthermore, GSCs express similar markers to NSCs including NESTIN, MUSASHI1, ZEB1, and SOX2 ^18, 19^.

The transcription factor ZEB1 is a key regulator of cancer stemness, tumor invasion and therapy resistance in GBM ^13^. It also has been found that HIF1α-ZEB1 axis mediate hypoxia induced mesenchymal shift in glioblastoma cells and invasive phenotype. Its presence results in enhanced stemness, tumorigenicity and lowered patient survival ^20, 21^. It has been found that patients with no ZEB1 expression showed greater survival and good response to TMZ chemotherapy. Another well-established transcriptional regulator of GSCs is SOX2. It has been found that SOX2 is induced by the ZEB1-miR-200 feedback loop in glioblastoma ^13^. SOX2 is found to be overexpressed in 90% of GBM patient samples and in the undifferentiated GSC populations, suggesting that it plays an important role in maintaining GSC stemness. Moreover, SOX2 silencing has been found to cause proliferation impairment, reduction in tumor growth and invasion and cell cycle arrest ^22^.

Many studies have established the critical role of receptor tyrosine kinases (RTKs) in GBM, including fibroblast growth factor receptors (FGFRs). For example, overexpression of FGFR1, specifically in the form of FGFR1-β, has been detected in GBM compared to normal levels of the receptor in the white matter ^23^. FGFR1 expression increases with WHO grade in astrocytomas, and it has been used as a marker for poor prognosis ^24^. It has been found that FGFR1 expression promotes tumor growth and invasion via AKT/MAPK and RAC1/CDC42 pathways, respectively ^25^. Moreover, FGFR1 point mutations (N546K and R576W) in the tyrosine kinase domain contribute to the growth of glioblastoma, as they result in a change in the surface charge of the protein enhancing protein-protein interactions and therefore the likelihood of FGFR1 autophosphorylation ^26^. By contrast, FGFR2 expression in the white matter is abundant, compared to malignant astrocytomas in which it is barely detectable ^23^. Also, *FGFR2* is considered a GBM-associated tumor suppressor gene and high expression of FGFR2 in GBM is associated with increased patient survival ^27, 28^. Low FGFR2 expression in GBM is attributed to the loss of heterozygosity of chromosome 10 found in 80% of GBMs, where the FGFR2 gene is localized (10q26) ^28, 29^.

Others and we have demonstrated that FGFR1 is expressed on GSCs, with FGF2 binding to FGFR1 activating downstream signaling pathways that maintain cancer stemness. Furthermore, the transcription factor ZEB1 regulates FGFR1 expression, thereby closing a positive feedback loop ^30-32^.

Here, we investigate the regional expression of FGFRs within the GBM tumor microenvironment and their association with GSCs. We quantify relative expression levels of FGFR1 and FGFR2 in GBM patient-derived xenograft models and compare expression within the tumor core and the invasion front.

## Materials and Methods

### Cell lines

Primary human glioblastoma cell lines L0, L1, and L2 were cultured as described previously ^32^. Briefly, cells were expanded in N2 medium containing 20 ng/ml EGF and 2% bovine serum albumin, passaged every 7 days and plated at 50,000 cells/ml. Lentiviral knockdown of FGFR1 and FGFR2 was performed as described ^32^ and confirmed by Western blot.

### Xenograft models

All animal experiments were carried out in accordance with UK Home Office regulations and the Animals (Scientific Procedures) Act 1986 (Home Office license PPL 30/3331). Mice were group-housed in 12-hour light/dark cycles in filter top cages with access to food and water *ad libitum*. Cages were cleaned weekly, and nesting material as well as plastic tunnels were provided for environmental enrichment. Intracranial implantation of GBM cells as performed as described in ^32^. Briefly, human GBM L2 cells ^13^ transduced with control, FGFR1 or FGFR2 knockdown lentiviral vectors carrying a GFP reporter were FACS purified prior to implantation. 50,000 cells were delivered in a total volume of 5 µl into the right hemisphere of female adult immunocompromised mice (n=3 per group) ^32^. Tumors were allowed to grow until the animals reached defined endpoint criteria, including body weight loss and/or neurological symptoms. Mice at endpoint were euthanized and transcardially perfused, the brains harvested, postfixed, embedded in optimal cutting temperature medium (OCT) and frozen. 30μm coronal sections were cut on a cryostat and used for immunostaining.

### Western blot

Protein lysates were extracted from control and FGFR1 and FGFR2 knockdown cells using RIPA buffer ^32^. The lysates were centrifuged, and a Bradford assay was used to determine protein concentration. Protein lysates were diluted with Laemmli sample buffer (Sigma Aldrich) and distilled water. The mixture was then heated at 55°C for 5 minutes. Proteins were separated using sodium dodecyl sulfate polyacrylamide gel electrophoresis (BioRad). Following gel electrophoresis, samples were transferred onto PVDF membranes (activated in methanol for 1 minute) using the Trans-Blot® TurboTM Transfer System (BioRad). Transferred membranes were washed in tris-buffered saline (TBS) for 10 minutes. The membrane was then blocked in 5% non-fat milk in tris-buffered saline-Tween (TBS-T) for 1 hour. The membrane was then washed twice in TBS-T for 10 minutes. The primary antibodies were diluted in antibody dilution or blocking solution (**Table S1**) and the membrane was incubated on a shaker in the primary antibody solution overnight at 4°C. The membrane was then washed in 3x in TBS-T for 5 minutes each. The membrane was then incubated with secondary antibody on a shaker for 1 hour. Antibody binding was visualized using chemiluminescence (BioRad) on a BioRad ChemiDocTM Imaging System.

### Immunofluorescence staining

Tissue sections were washed twice for 10 minutes each in PBS-T (PBS containing 0.1% triton X-100). Sections were then blocked in FSB-T (Fish Skin Gelatine Blocking Buffer Triton X-100; 500ml PBS/TBS, 5g BSA (1%), 2ml Teleostean gelatin (0.2%), 500μl Triton X-100, 0.1g Sodium Azide) for 1 hour at room temperature on a shaker. Sections were incubated overnight at 4 °C in primary antibodies diluted in FSB-T (**Table S1**). Sections were washed 3x for 10 minutes each in PBS-T then incubated with fluorescence-labelled secondary antibodies diluted in FSB-T for 3 hours at room temperature in the dark on a shaker. Sections were incubated with Hoechst 33342 nuclear staining solution (Thermo Fisher Scientific) diluted in PBS-T for 5 minutes at room temperature in the dark on a shaker. Sections were then washed 3x for 10 minutes each in PBS-T before mounting on microscopic slides, and coverslipping with ProLongTM Diamond Antifade Mountant (Invitrogen). Slides are kept in the dark at 4 °C prior to imaging.

### Microscopy and image processing

Slides were visualized using a confocal microscope (Zeiss LSM710). Images were obtained using ZEN software. Images were processed and the fluorescence signals were quantified using ImageJ software (https://imagej.nih.gov/ij/). The maximum intensity projection of the core and the invasion front of the tumor were measured using ImageJ.

### RNA sequencing and data processing

Total RNA was extracted from human glioblastoma cells using the RNeasy kit according to the manufacturer’s instructions (Qiagen). RNA quality was assessed on a Bioanalyzer (Agilent) with all RNA integrity (RIN) values 8 or greater. Library preparation for RNA sequencing was performed using the Ion Total RNA-Seq V2 kit and Ion Xpress RNA-Seq Barcode kit (Thermo Fisher). Library quality control was performed using the Ion Library Taqan Quantitation kit (Thermo Fisher). RNA sequencing was run on an IonChef System using the Ion PI HiQ Chef kit and Ion PI chips (Thermo Fisher).

Trimming and quality control of raw sequencing data was performed using FastQC, followed by read mapping using STAR ^33^. The read depth was at least 10 million mapped reads per sample. Raw read counts and RPKM values were calculated for individual exons and transcripts using an in-house script at Wales Gene Park. Differentially expressed genes were identified using DEseq2 ^34^. The resultant p-values were corrected for multiple testing and false discovery issues using the FDR method ^35^.

### Ingenuity pathway analysis

Data were analyzed using QIAGEN Ingenuity Pathway Analysis. Data sets containing gene identifiers and corresponding data measurement values were uploaded into the application. Each identifier was mapped to its corresponding object in QIAGEN’s Knowledge Base. An expression p value cutoff of 0.05 was set to identify molecules whose expression (or phosphorylation) was significantly perturbed. These molecules, called Network Eligible molecules, were overlaid onto a global molecular network developed from information contained in the QIAGEN Knowledge Base. Networks of Network Eligible Molecules were then algorithmically generated based on their connectivity.

Canonical pathways analysis identified the pathways from the QIAGEN Ingenuity Pathway Analysis library of canonical pathways that were most significant to the data set. Molecules from the data set that met the expression p value cutoff of 0.05 and were associated with a canonical pathway in the QIAGEN Knowledge Base were considered for the analysis. The significance of the association between the data set and the canonical pathway was measured in two ways: 1) A ratio of the number of molecules from the data set that map to the pathway divided by the total number of molecules that map to the canonical pathway is displayed; and 2) A right-tailed Fisher’s Exact Test was used to calculate a p-value determining the probability that the association between the genes in the dataset and the canonical pathway is explained by chance alone.

The Diseases & Functions Analysis identified the biological functions that were most significant from the data set. Molecules from the dataset that met the expression p value cutoff of 0.05 and were associated with biological functions in the QIAGEN Knowledge Base were considered for the analysis. A right-tailed Fisher’s Exact Test was used to calculate a p-value determining the probability that each biological function and/or disease assigned to that data set is due to chance alone.

The Functional Analysis of a network identified the biological functions that were most significant to the molecules in the network. The network molecules associated with biological functions in the QIAGEN Knowledge Base were considered for the analysis. A right-tailed Fisher’s Exact Test was used to calculate a p-value determining the probability that each biological function and/or disease assigned to that network is due to chance alone.

### Statistical analysis

Statistical analyses were performed using IBM SPSS® software. The Mann-Whitney U test was applied as the data were not normally distributed. A p-value less than 0.05 was considered significant.

## Results

FGFR1 and FGFR2 show an inverse relationship with GBM malignancy. Whereas high expression of FGFR1 is associated with poorer patient survival, higher expression of FGFR2 conveys a better prognosis ^24, 28^. Nevertheless, we have previously found that primary patient-derived GBM cells express FGFR2 *in vitro* ^32^. Therefore, we wanted to elucidate if FGFR1 and FGFR2 are expressed in GBM *in vivo* and if there are receptor-specific patterns of expression, specifically comparing tumor core and invasive areas. We chose to determine FGFR protein expression in patient-derived xenografts, as it is easier to visualize the core of the tumor mass as well as areas of tumor invasion within the same tissue section in these samples.

### Validation of anti-FGFR antibodies

To validate the specificity of anti-FGFR antibodies, we performed Western blot analysis of GBM cells transduced with scramble control or FGFR knockdown constructs (**Fig. 1**). After testing 2 different monoclonal antibodies against FGFR1 (clone M17A3, raised in mouse and D8E4, raised in rabbit), we excluded clone M17A3 as there were no bands visible on the Western blot (**Fig. 1A, B**). The antibody against FGFR2 produced clearly visible bands and was used for further experiments (**Fig. 1C**). Western blots showed that transduction of GBM cells with either shFGFR1 or shFGFR2 constructs ablated the targeted receptor, compared to GBM cells transduced with non-targeting constructs. Next, we used immunofluorescence staining of orthotopic xenografts of control, as well as FGFR1 and FGFR2 knockdown cells. We validated absence of FGFR1 immunostaining in xenografts of shFGFR1 cells compared to controls, as well as lack of FGFR2 staining in shFGFR2 xenografts (**Fig. 1D, E**). In both cases, we used human-specific anti-Vimentin or anti-Nestin antibodies to label xenografted GBM cells.

**Figure 1:**
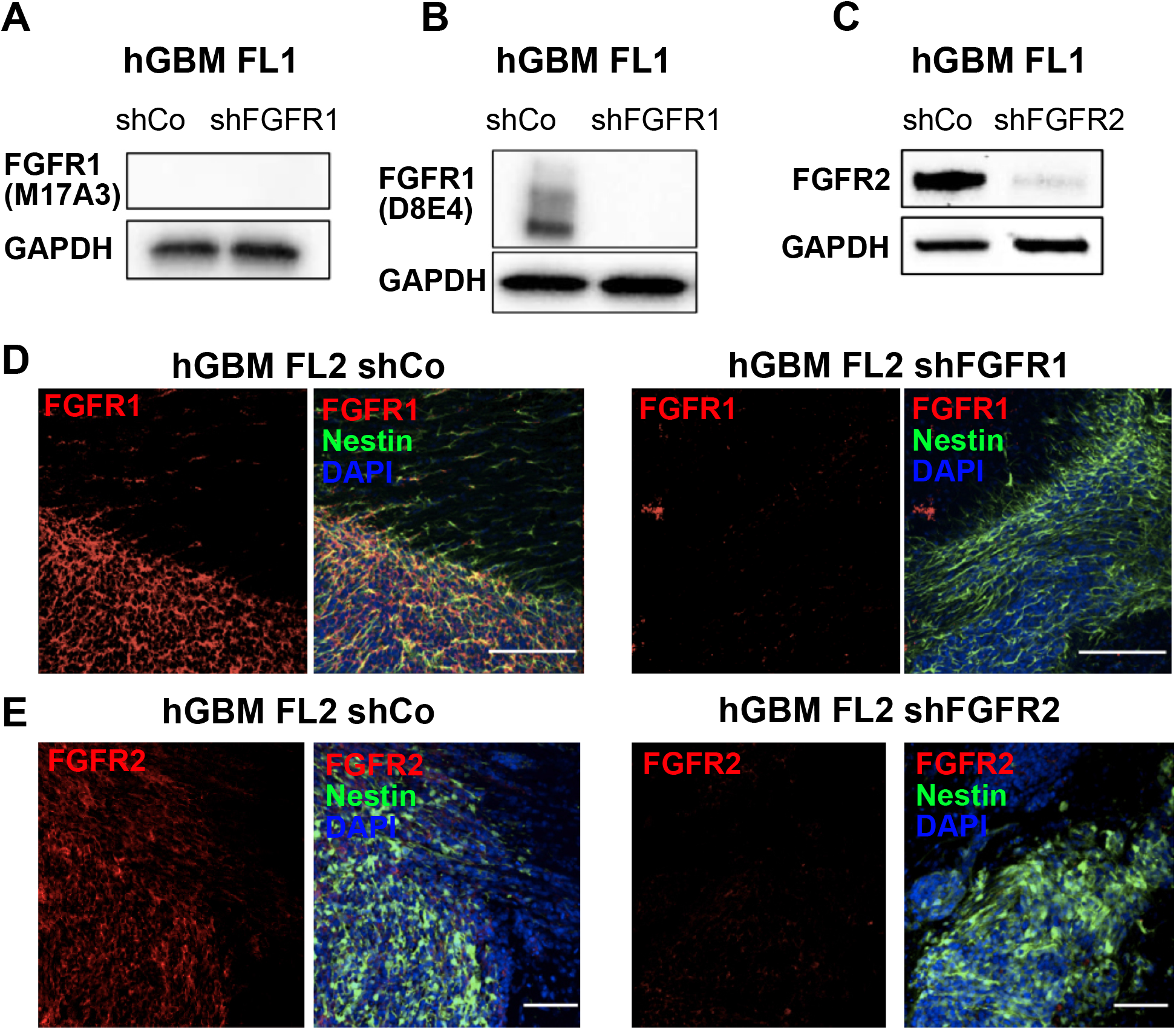
Expression of FGFR1 and FGFR2 in GBM xenografts. **(A)** A mouse monoclonal antibody against FGFR1 (M17A3) yielded no signal in Western blot and was not further investigated in this study. **(B)** Rabbit monoclonal antibody (D8E4) showed FGFR1-specific bands which are not present in FGFR1 knockdown cells (shFGFR1). **(C)** Mouse monoclonal antibody (D4L2V) against FGFR2 showed FGFR2-specific bands which are not present in FGFR2 knockdown cells (shFGFR2). GAPDH was used as loading control in A-C. **(D)** FGFR1 immunofluorescence staining of GBM xenografts shows a specific signal that overlaps with human-specific Nestin in tumor cells, while FGFR1 knockdown cells show no signal. Scale bar 80 μm. **(E)**. FGFR2 immunofluorescence staining of GBM xenografts shows a specific signal that overlaps with human specific Nestin in tumor cells, while FGFR2 knockdown cells show no signal. Scale bar 50 μm.

### FGFR1 and FGFR2 show differential spatial distribution in xenografted GBM

After validating anti-FGFR antibodies, we determined expression of FGFR1 in orthotopic patient-derived xenograft models of GBM, comparing central areas of the tumor core to the invasive edge. We used coronal sections from three recipient mice containing xenografted tumors and quantified mean fluorescence intensities across multiple areas (n>14 per brain) that were either classified as tumor core (containing a mass of tumor cells) or invasive edge (containing infiltrating tumor cells intermixed with host cells). We also quantified mean fluorescence intensity from sections of mice xenografted with FGFR1 knockdown tumors as negative controls. There was no significant difference between fluorescence intensities in the tumor core and the invasive edge, confirming visual observations that FGFR1 is expressed throughout the tumor (**Fig.2 A, B)**. Decreased FGFR1 expression in FGFR1 knockdown cells resulted in lower fluorescence intensity of FGFR1 staining in xenografted tumors, with no appreciable difference between tumor core and invasive edge (**Fig. 2C**).

**Figure 2:**
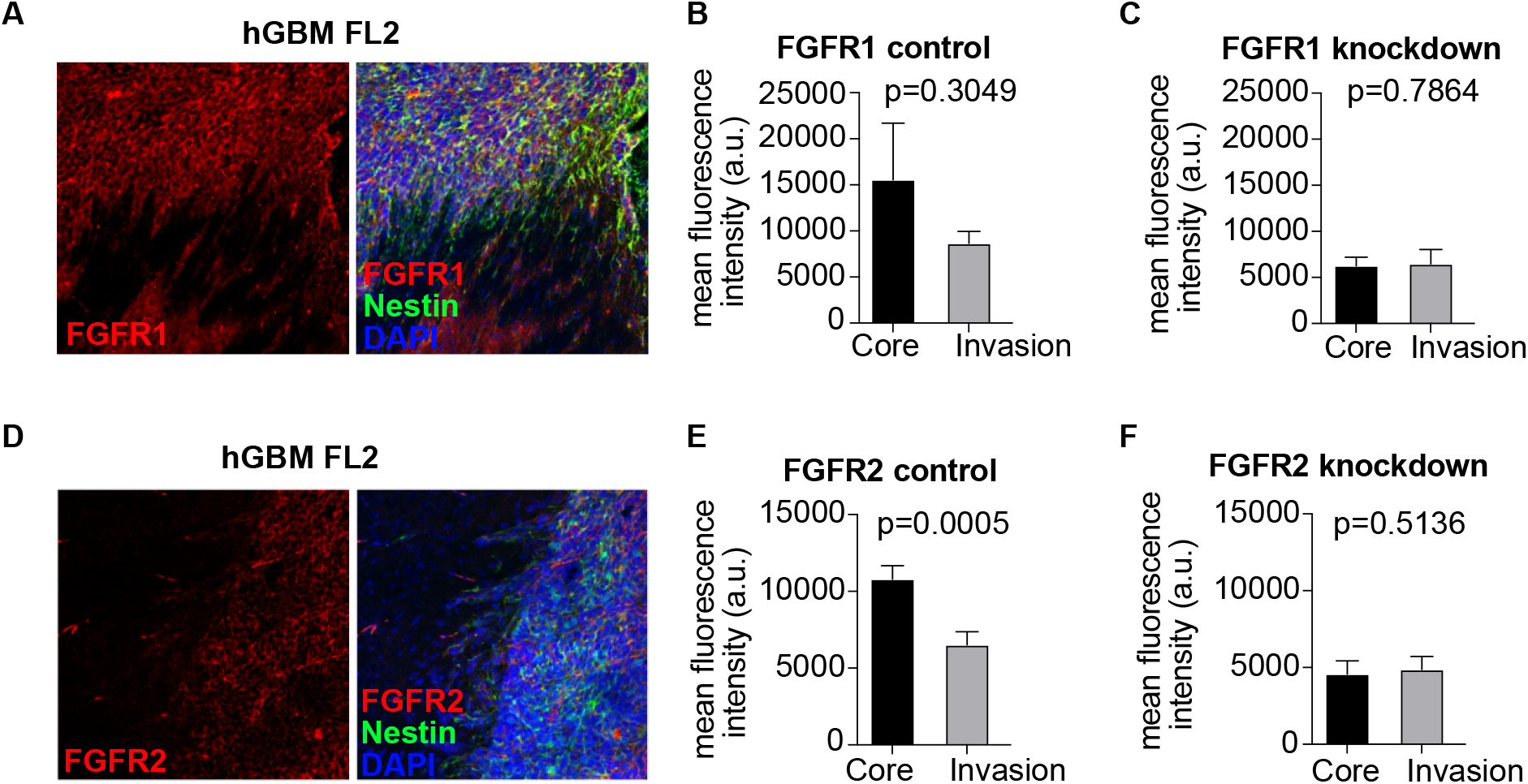
Spatial distribution of FGFR1 and FGFR2 in GBM xenografts. **(A)** GBM xenograft immunofluorescence staining shows expression FGFR1 in the tumor core (top part of image) and invasive cells (bottom part of image). **(B)** Quantification of mean fluorescence intensity for FGFR1 in tumor core and invasive areas shows no significant difference (n=3 animals). **(C)** Quantification of mean fluorescence intensity for FGFR1 in xenografted FGFR1 knockdown cells shows that the fluorescence signal is lower than in control cells and no difference between tumor areas (n=3 animals). **(D)** Immunofluorescence staining for FGFR1 in GBM xenografts shows strong staining in the tumor core (right part of image), which is absent in invasive cells (left part of image). **(E)** Quantification of mean fluorescence intensity for FGFR2 shows significant difference between tumor core and invasive areas (n=3 animals). **(F)** Quantification of mean fluorescence intensity for FGFR2 in xenografted FGFR2 knockdown cells shows that the fluorescence signal is lower than in control cells and no difference between tumor areas (n=3 animals). Scale bar 80 μm.

We next evaluated the expression patterns of FGFR2 in GBM xenografts, again focusing on potential differences between the tumor core and the invasive edge. We quantified mean fluorescence intensities of xenografted tumors (n>14 areas per brain), using xenografted FGFR2 knockdown GBM cells as negative controls. We found that the distribution of FGFR2 shows spatial heterogeneity, with high expression in tumor core areas, and low expression in invasive areas (**Fig. 2D, E**). As expected, ablation of FGFR2 resulted in low fluorescence intensity of FGFR2 staining (**Fig. 2F**). Thus, FGFR1 and FGFR2 show differential distribution within xenografted human GBM, with FGFR1 expressed in invasive areas, while FGFR2 at background levels in invasive cells.

### Knockdown of FGFR1 reduces tumor invasion, but FGFR2 knockdown does not

Because invasive GBM cells show differential expression of FGFR1 and FGFR2, we asked whether FGFR1 is functionally relevant for invasive GBM cells. To determine whether FGFR1 is functionally relevant for GBM cell migration/invasion we performed sphere outgrowth assays *in vitro* and quantified tumor invasion *in vivo*. Knockdown of FGFR1 in human patient derived GBM cell lines resulted in reduced cell migration compared to non-targeting controls (**Fig. 3A**). Likewise, ablation of FGFR1 resulted in a profound reduction in tumor invasion in xenografted GBM tumors derived from FGFR1 knockdown cells or non-targeting controls (**Fig. 3B, C**). These results are in line with our previous work demonstrating functional relevance of FGFR1 for stemness in GBM and increased invasive propensity of GBM cancer stem cells ^13, 32, 36^. Hence, FGFR1 is functionally relevant for tumor invasion in GBM.

**Figure 3:**
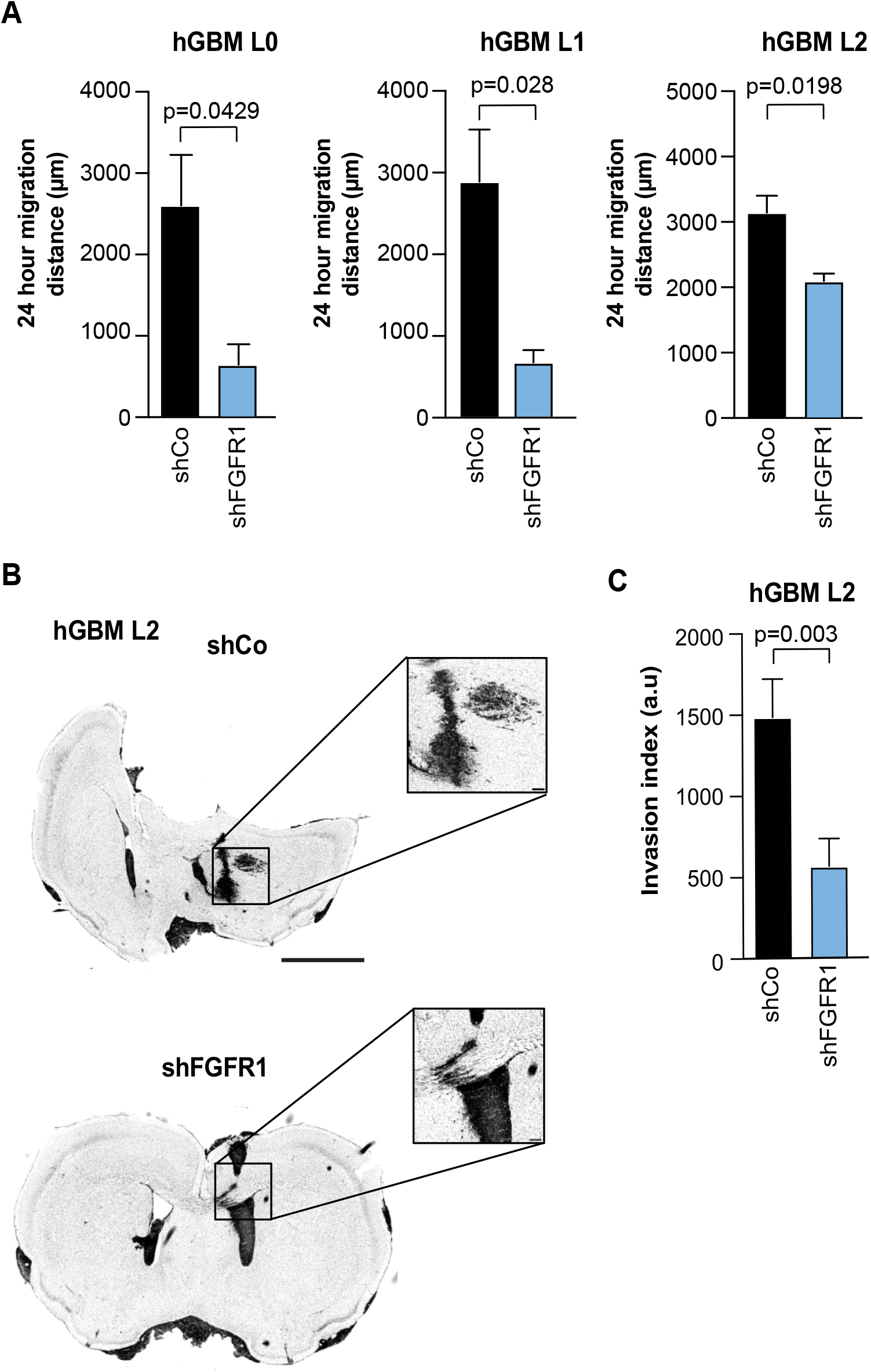
FGFR1 regulates GBM cell migration and tumor invasion. **(A)** Quantification of cell migration distances (sphere outgrowth assay) for three different primary patient derived GBM cell lines (n=3 independent experiments each). FGFR1 knockdown (shFGFR1) significantly reduced cell migration compared to non-targeting controls (shCo). **(B)** Representative images (inverted grayscale of nuclear stain) of orthotopic xenografts of hGBM L2 expressing non-targeting control (shCo) or FGFR1 knockdown (shFGFR1) vectors. Insets show magnification of boxed areas. Tumors derived from control cells are diffusely invasive while shFGFR1-derived tumors are less invasive. Scale bar 5 mm (insets 50 µm). **(C)** Quantification of tumor invasion of hGBM L2 xenografts (n=3 animals) shows a significant reduction in tumor invasion in shFGFR1-derived tumors.

Because FGFR2 is not expressed in areas of GBM invasion, we speculated that FGFR2 is not regulating tumor invasion in GBM. To test this, we compared cell migration in sphere outgrowth assays *in vitro* and tumor invasion in xenografts *in vivo*. FGFR2 knockdown did not reduce cell migration compared to non-targeting controls (**Fig. 4A**). Contrastingly to loss of FGFR1, which decreased tumor invasion, xenotransplanted FGFR2 knockdown cells showed comparable tumor invasion as non-targeted control cells (**Fig. 4B, C**). As expected from its expression pattern, FGFR2 loss is not affecting invasive GBM cells.

**Figure 4:**
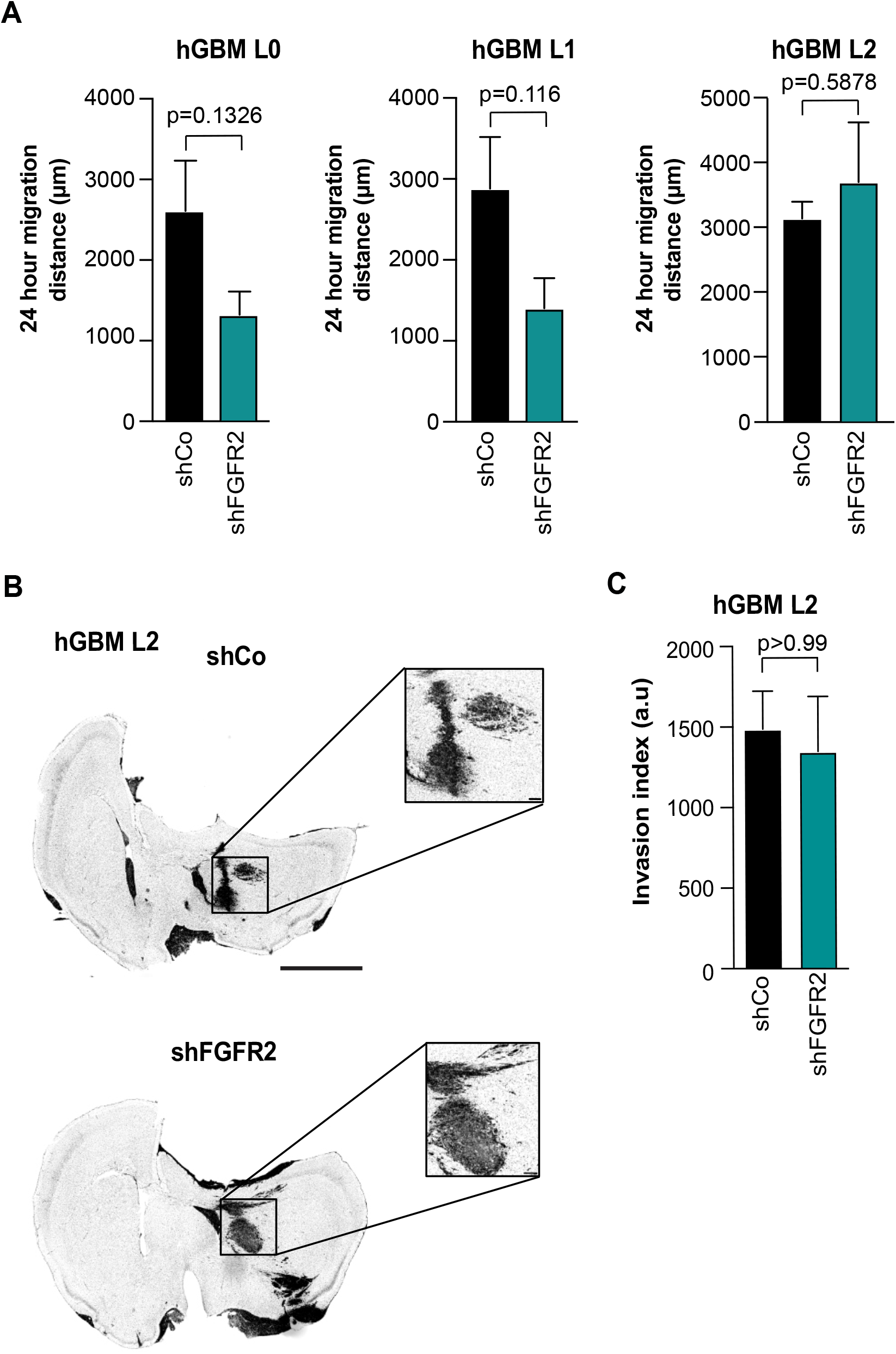
FGFR2 does not affect GBM cell migration or tumor invasion. **(A)** Quantification of cell migration distances (sphere outgrowth assay) for three different primary patient derived GBM cell lines (n=3 independent experiments each). Observed changes in cell migration following FGFR2 knockdown (shFGFR2) were not significant. **(B)** Representative images (inverted grayscale of nuclear stain) of orthotopic xenografts of hGBM L2 expressing control (shCo) or FGFR2 knockdown (shFGFR2) vectors. Insets show magnification of boxed areas. Tumors derived from control and shFGFR2 cells are diffusely invasive. Scale bar 5 mm (insets 50 µm). **(C)** Quantification of tumor invasion of hGBM L2 xenografts (n=3 animals) shows no change in tumor invasion in shFGFR2-derived tumors.

### Transcriptional profiling of FGFR1 and FGFR2 knockdown cells identifies gene networks associated with tumor invasion

To better understand the relevance of FGFR1 for GBM cell migration and invasion and to identify potential gene-regulatory networks involved in tumor invasion of FGFR1-expressing GBM cells, we performed transcriptional profiling using RNA sequencing. We compared knockdown of FGFR1 and FGFR2 to non-targeting controls across 3 primary patient-derived human GBM cell lines, using single-end RNA-sequencing (RNA-seq; **Fig. 5A**). All cells were treated with FGF2 for 48 hours prior to RNA preparation. Differentially expressed genes were identified for FGFR1 knockdown versus control and FGFR2 knockdown versus control using DEseq2 ^34^. We used gene set enrichment analysis (GSEA) to identify relevant gene sets in FGFR1 and FGFR2 knockdown samples. This revealed that gene sets for epithelial-mesenchymal transition were significantly enriched in controls compared to FGFR1 knockdown (**Fig. 5B**), but not FGFR2 knockdown (not shown), validating our previously reported link between FGFR1 and the transcription factor ZEB1 ^32^.

**Figure 5:**
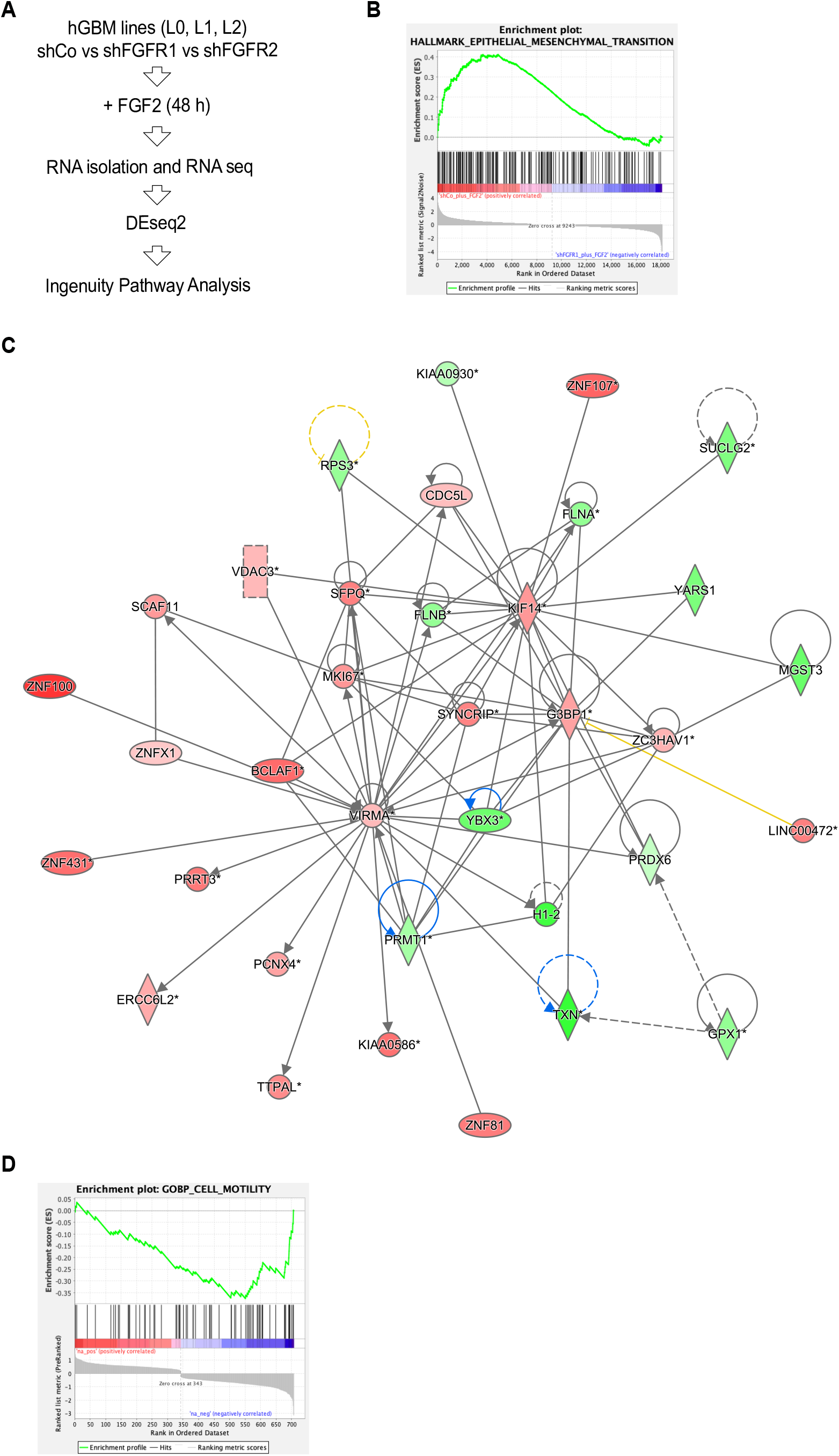
RNA-seq analysis of control, FGFR1- and FGFR2 knockdown GBM cells. **(A)** Schematic of experimental design. Three primary patient derived GBM cell lines were transduced with non-targeting control (shCo) or knockdown vectors for FGFR1 (shFGFR1) or FGFR2 (shFGFR2). Cells were treated with FGF2 for 48 hours prior to RNA isolation and library preparation. DEseq2 was used to identify differentially expressed genes in RNA-seq datasets, which were subsequently analyzed by Ingenuity Pathway Analysis. **(B)** GSEA plot shows enrichment of epithelial-mesenchymal transition signature in shCo compared to shFGFR1 cells (NES=1.22, p<0.0001). **(C)** Ingenuity pathways map of cell motility-associated gene network that is regulated by FGFR1. Green indicates decreased expression and red indicates increased expression of genes. Arrows indicate regulatory relationships. Direct interactions are indicated by solid lines, indirect interactions by dashed lines. **(D)** GSEA plot shows enrichment for cell motility associated genes in shCo compared to shFGFR1 cells (NES=-1.97, p<0.0001).

We then used QIAGEN Ingenuity Pathway Analysis ^37^ for a comparative analysis of FGFR1 knockdown versus FGFR2 knockdown results. IPA analysis yielded a total of 1564 differentially expressed genes in FGFR1 knockdown cells and 1851 differentially expressed genes in FGFR2 knockdown cells that passed significance threshold. IPA comparison between FGFR1 and FGFR2 knockdown revealed 56 differentially activated canonical pathways that reached the significance threshold of a Benjamini-Hochberg p value of 0.05 in FGFR1 knockdown cells. Of note, the network ‘RhoA signaling’ showed a negative z-score in FGFR1 knockdown cells (meaning network genes are more likely downregulated) but a positive z-score in FGFR2 knockdown cells (Benjamini-Hochberg p-value 0.006). RhoA signaling is linked to cell shape, locomotion, and polarity, and downregulation of RhoA network genes following loss of FGFR1 supports decreased cell motility and invasion in these cells. A comparison of biological functions in the Qiagen knowledge base showed significant association of ‘Cell movement of tumor cell lines’ and ‘Migration of tumor cell lines’ (both B-H p value 0.003) with FGFR1 knockdown, but not FGFR2 knockdown cells. We used Qiagen’s functional network analysis to further identify gene networks associated with cell motility and tumor invasion, which identified a network of cell motility-associated gene targets (**Fig. 5C** and **Table 1**).

**Table 1:**
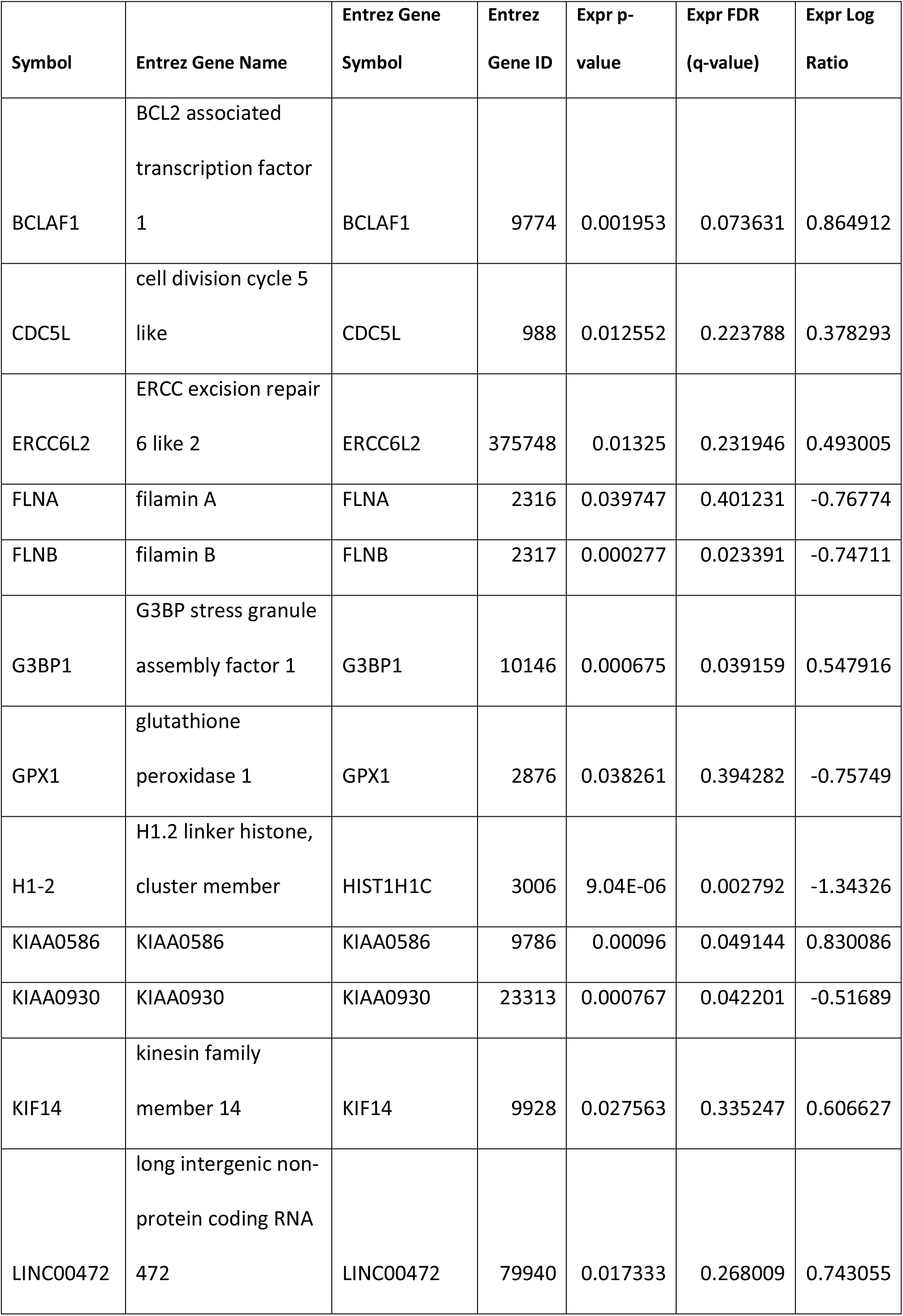
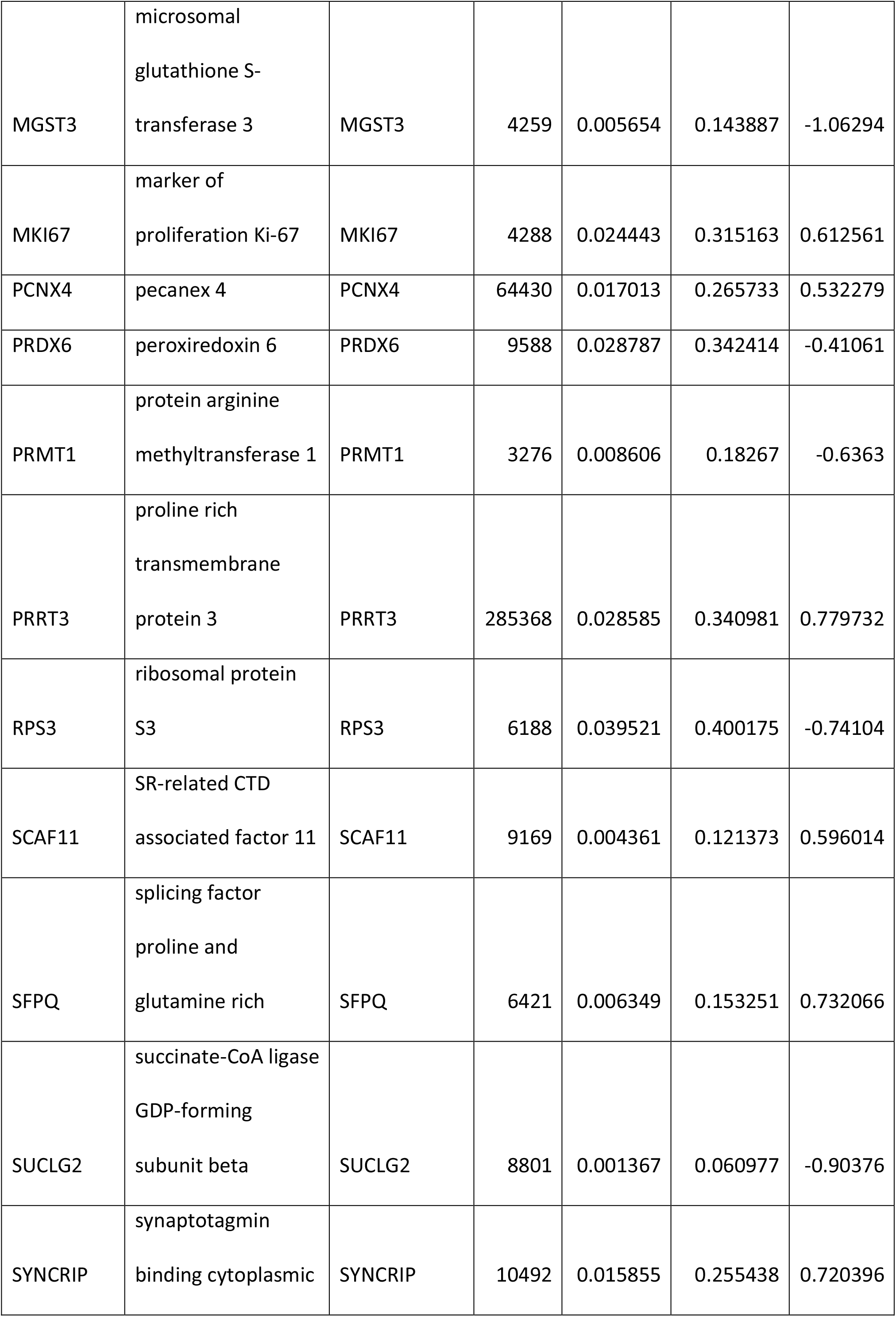
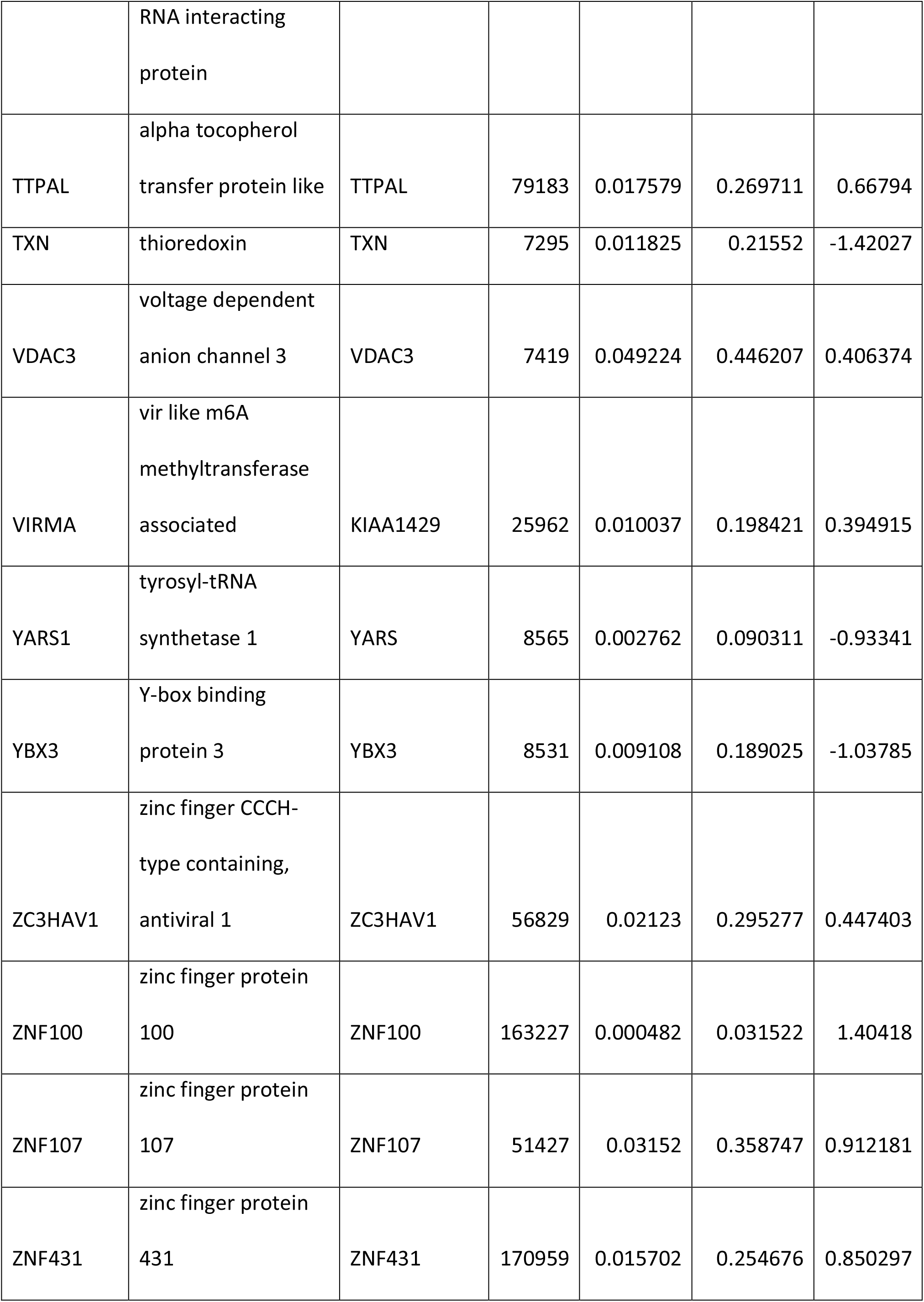
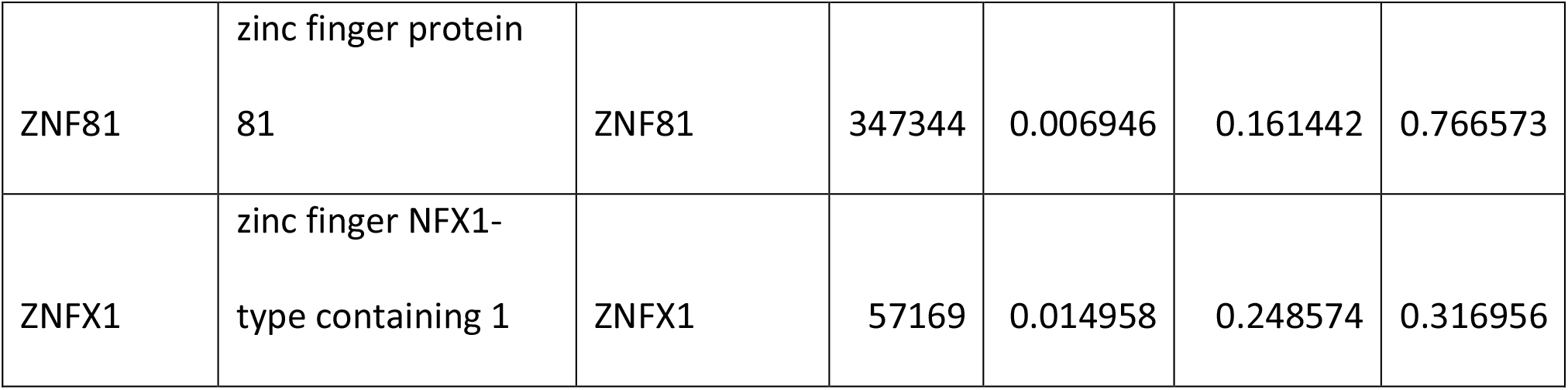
Cell motility-associated gene targets regulated by FGFR1. Table lists candidates from Ingenuity Pathway Analysis with expression p-values and log ratio.

To identify potential candidate gene sets involved in cell migration that are specific to FGFR1 signaling, we curated a list of differentially expressed genes from FGFR1 knockdown cells that are not changed following FGFR2 knockdown. We identified 715 differentially expressed genes that are specifically regulated by FGFR1. We ranked these 715 candidates by expression log fold change and performed gene set enrichment analysis for gene ontologies associated with cell motility. This revealed a significantly lower expression of candidate genes linked associated with GO terms ‘cell motility’, ‘wound healing’, ‘cell adhesion’, ‘taxis’, ‘locomotion’, ‘amoeboidal type cell migration’, and ‘regulation of locomotion’ (FDR q-values 0.021, 0.020, 0.024, 0.033, 0.040, 0.049, 0.049, respectively). An exemplary enrichment plot for ‘cell motility’ is shown in **Fig. 5D** and candidates are listed in **Table 2**. Enrichment plots for other GO terms are included as supplementary data (**Fig. S1)**.

**Table 2:** Gene set enrichment analysis candidates. Table lists candidates from GSEA associated with cell motility with rank and running enrichment score.

| **NAME** | **SYMBOL** | **RANK**<br>**IN**<br>**GENE**<br>**LIST** | **RANK**<br>**METRIC**<br>**SCORE** | **RUNNING**<br>**ES** | **CORE**<br>**ENRICHMENT** |
| --- | --- | --- | --- | --- | --- |
| row_0 | TP53INP1 | 3 | 1.27764904 | 0.01921398 | No |
| row_1 | CDKN2B-AS1 | 7 | 1.13283801 | 0.03571892 | No |
| row_2 | ASPM | 41 | 0.83374202 | -2.46E-04 | No |
| row_3 | PARVA | 67 | 0.73762798 | -0.0255098 | No |
| row_4 | DEPDC1B | 117 | 0.62990201 | -0.0902885 | No |
| row_5 | PARD6B | 126 | 0.615991 | -0.0912649 | No |
| row_6 | KIF14 | 132 | 0.60662699 | -0.087729 | No |
| row_7 | JAGN1 | 141 | 0.59868002 | -0.0890293 | No |
| row_8 | TBC1D24 | 145 | 0.59428799 | -0.0825992 | No |
| row_9 | BRAF | 173 | 0.559798 | -0.1143144 | No |
| row_10 | SYDE1 | 175 | 0.55852997 | -0.1054282 | No |
| row_11 | MAZ | 178 | 0.55233002 | -0.0982206 | No |
| row_12 | PDCD6 | 198 | 0.53384399 | -0.1179212 | No |
| row_13 | ANLN | 210 | 0.51605499 | -0.1254547 | No |
| row_14 | ARPIN | 226 | 0.49285701 | -0.1396721 | No |
| row_15 | KANK2 | 228 | 0.49211401 | -0.1320285 | No |
| row_16 | PHACTR4 | 243 | 0.47075301 | -0.1450969 | No |
| row_17 | TAOK2 | 256 | 0.445806 | -0.155507 | No |
| row_18 | CEP131 | 259 | 0.43920201 | -0.1504157 | No |
| row_19 | CRK | 280 | 0.40059701 | -0.1741716 | No |
| row_20 | DIAPH1 | 327 | 0.30548501 | -0.2403318 | No |
| row_21 | STK4 | 330 | 0.30057299 | -0.2378338 | No |
| row_22 | RAF1 | 336 | 0.26874399 | -0.2406188 | No |
| row_23 | RHOA | 339 | 0.25658399 | -0.2389438 | No |
| row_24 | AMOTL1 | 340 | 0.20852999 | -0.2350428 | No |
| row_25 | ARRB2 | 351 | -0.30765 | -0.2449124 | No |
| row_26 | BSG | 360 | -0.35071 | -0.2508516 | No |
| row_27 | STK24 | 364 | -0.36667 | -0.2486796 | No |
| row_28 | CDC42BPB | 382 | -0.40257 | -0.2677111 | No |
| row_29 | PLXNA1 | 383 | -0.40391 | -0.260155 | No |
| row_30 | MYO18A | 390 | -0.4181 | -0.2617084 | No |
| row_31 | LGMN | 418 | -0.48845 | -0.2947583 | No |
| row_32 | CD151 | 424 | -0.49881 | -0.2932394 | No |
| row_33 | ARPC3 | 435 | -0.5158 | -0.2992151 | No |
| row_34 | FUT10 | 438 | -0.52039 | -0.292605 | No |
| row_35 | SEMA6C | 442 | -0.52626 | -0.2874475 | No |
| row_36 | IGSF8 | 466 | -0.57275 | -0.3126704 | No |
| row_37 | PRKCE | 505 | -0.64769 | -0.3599288 | No |
| row_38 | ADD2 | 506 | -0.64852 | -0.3477967 | No |
| row_39 | CD99 | 516 | -0.66842 | -0.3493547 | No |
| row_40 | MITF | 520 | -0.68504 | -0.3412269 | No |
| row_41 | SPHK1 | 537 | -0.72581 | -0.3526489 | No |
| row_42 | FLNA | 552 | -0.76774 | -0.3601615 | Yes |
| row_43 | PTPRF | 557 | -0.77338 | -0.3519436 | Yes |
| row_44 | DNAH1 | 561 | -0.78178 | -0.342006 | Yes |
| row_45 | MERTK | 565 | -0.79203 | -0.3318767 | Yes |
| row_46 | NINJ1 | 569 | -0.80225 | -0.3215562 | Yes |
| row_47 | CSNK2B | 575 | -0.82087 | -0.3140124 | Yes |
| row_48 | KCTD13 | 576 | -0.82126 | -0.2986487 | Yes |
| row_49 | MDK | 589 | -0.86355 | -0.301244 | Yes |
| row_50 | SEMA4C | 590 | -0.86373 | -0.2850859 | Yes |
| row_51 | TTC21A | 595 | -0.87663 | -0.2749364 | Yes |
| row_52 | MAPK15 | 596 | -0.87707 | -0.2585287 | Yes |
| row_53 | PLCG2 | 600 | -0.88218 | -0.2467129 | Yes |
| row_54 | PTP4A3 | 605 | -0.89508 | -0.2362183 | Yes |
| row_55 | SERPINF1 | 607 | -0.90402 | -0.220869 | Yes |
| row_56 | LAMC3 | 630 | -0.97518 | -0.2370009 | Yes |
| row_57 | SERPINE2 | 649 | -1.07105 | -0.2450893 | Yes |
| row_58 | SDC4 | 677 | -1.31201 | -0.2627325 | Yes |
| row_59 | HBEGF | 679 | -1.31889 | -0.2396221 | Yes |
| row_60 | GADD45A | 681 | -1.3538899 | -0.2158568 | Yes |
| row_61 | TRIB1 | 691 | -1.52238 | -0.2014396 | Yes |
| row_62 | DDIT4 | 692 | -1.5403399 | -0.1726238 | Yes |
| row_63 | CARMIL3 | 693 | -1.56016 | -0.1434373 | Yes |
| row_64 | FGFR1 | 694 | -1.58595 | -0.1137683 | Yes |
| row_65 | PLAU | 699 | -1.79008 | -0.0865306 | Yes |
| row_66 | DDIT3 | 704 | -2.0682001 | -0.05409 | Yes |
| row_67 | SERPINE1 | 706 | -3.0584099 | 0.00156241 | Yes |

Together, our results strongly support that FGFR1, but not FGFR2, promotes tumor invasion in glioblastoma, linking spatial distribution of receptor expression within glioblastoma to function on cell motility. We identify relevant gene sets associated with cell motility that are regulated by FGFR1.

## Discussion

Among the four main FGFRs, expression levels of FGFR1 and FGFR2 in glioblastoma show a significant but divergent association with patient survival. High expression levels of FGFR1 are linked to poor survival and FGFR1 is a regulator of cancer stem cells and therapy resistance in glioblastoma ^30-32^. By contrast, the functions of FGFR2 in glioblastoma remain incompletely understood. High expression levels of FGFR2 are associated with increased survival, and downregulation of FGFR2 is correlated with increased proliferation ^28^.

Here, we investigated the distribution and functional relevance of FGFR1 and FGFR2 in glioblastoma primary patient derived cell lines and xenografts. We found FGFR1 distributed across the tumor mass and diffusely infiltrating glioblastoma cells, while FGFR2 expression is confined to the tumor mass and notably absent in invasive cells. We previously reported that FGFR1 but not FGFR2 or FGFR3 regulated glioblastoma cell proliferation *in vitro* ^32^. Thus, the potential functions of FGFR2 within the tumor mass remain unclear and need further investigation. The differential expression of FGFR1/FGFR2 in invasive glioblastoma cells is reflected in the functional consequences of FGFR1 and FGFR2 knockdown, as only ablation of FGFR1, but not of FGFR2, reduced glioblastoma cell migration and tumor invasion.

FGFR1 expression and/or amplification has been linked to epithelial-mesenchymal transition and metastasis in other cancers, including lung cancer, bladder and urothelial cancer, and gastric cancer ^38-41^. Of note, a previous study demonstrated that isoform switching of FGFR1 promotes tumor invasion ^42^. We have not tested whether different FGFR1 isoforms are expressed in the tumor mass versus invasive cells, and this warrants further investigation. In glioblastoma, FGFR1 is known to regulate cancer stemness and therapy resistance through the transcription factor ZEB1 ^30-32^. We have previously shown that ZEB1 is important for glioblastoma invasion by regulating expression of the axon guidance molecule ROBO1 ^13^. To identify gene regulatory networks related to FGFR1-driven glioblastoma tumor invasion, we compared transcriptomes of control, FGFR1 knockdown, and FGFR2 knockdown glioblastoma cells using RNA-Seq. This revealed 715 transcripts which are specifically regulated by FGFR1 and not FGFR2 following stimulation with FGF2. Interestingly, ROBO1 expression was not altered in FGFR1 knockdown cells (data not shown), indicating that FGFR1-mediated glioblastoma invasion may rely on other effector pathways. To further investigate how FGFR1 regulates glioblastoma invasion, we used IPA to identify cell motility-associated gene regulatory networks that are dependent on FGFR1. This revealed a network consisting of 35 candidates and the key nodes VIRMA, KIF14, and G3BP1, all of which are significantly higher connected within this network. Hence, these nodes may constitute relevant targets to disrupt glioblastoma invasiveness. To our knowledge, these molecules have not been investigated in the context of glioblastoma invasion. Expression of KIF14 is enriched in female patient specimens ^43^ and KIF14 inhibition reduced cell growth and increased apoptosis in glioblastoma cell lines ^44^. Knockdown of G3BP1 sensitized glioma cell lines to chemotherapy ^45^.

In summary, our study demonstrates differential expression of FGFR1 and FGFR2 on invasive glioblastoma cells and provides functional and transcriptomic evidence that FGFR1 regulates tumor invasion in glioblastoma. We found candidate gene targets at the center of a regulator network associated with cell motility that warrant further investigation.

## Acknowledgements

The authors would like to thank Dr K. Ashelford from Wales Gene Park for assistance with processing of raw sequencing data. FAS is supported by MRC grant MR/S007709/1.

## Declaration

The authors declare no conflict of interest.

## Figure and Table Legends

**Figure S1:**
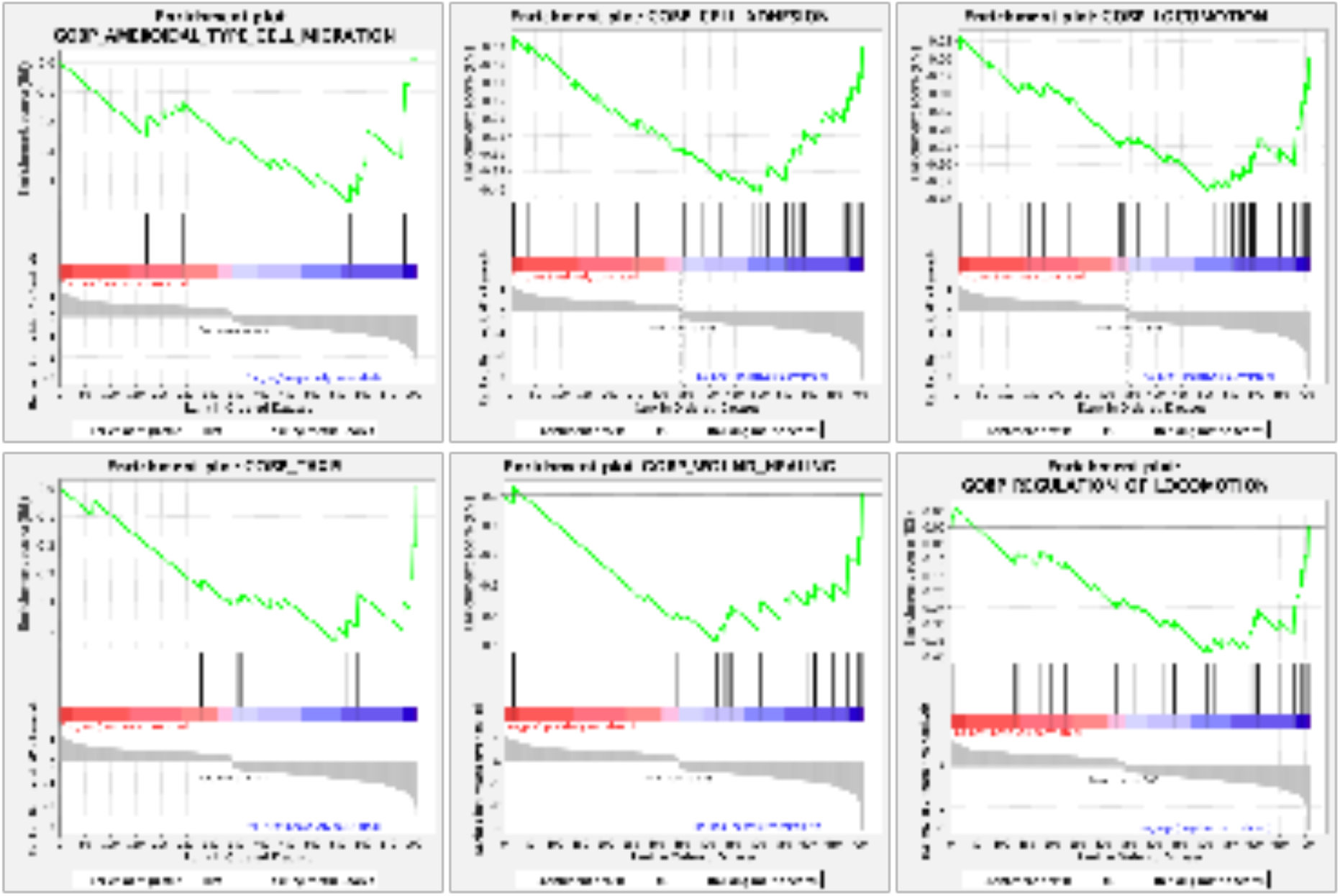
Gene set enrichment plots associated with FGFR1. Enrichment plots for GO terms ‘wound healing’, ‘cell adhesion’, ‘taxis’, ‘locomotion’, ‘amoeboidal type cell migration’, and ‘regulation of locomotion’ (NES= -1.96, -1.92, -1.85, -1.81, -1.81, -1.77, -1.77 respectively; p= 0.001, 0.001, 0.002, 0.015, 0.013, 0.006, respectively).

**Table S1:**
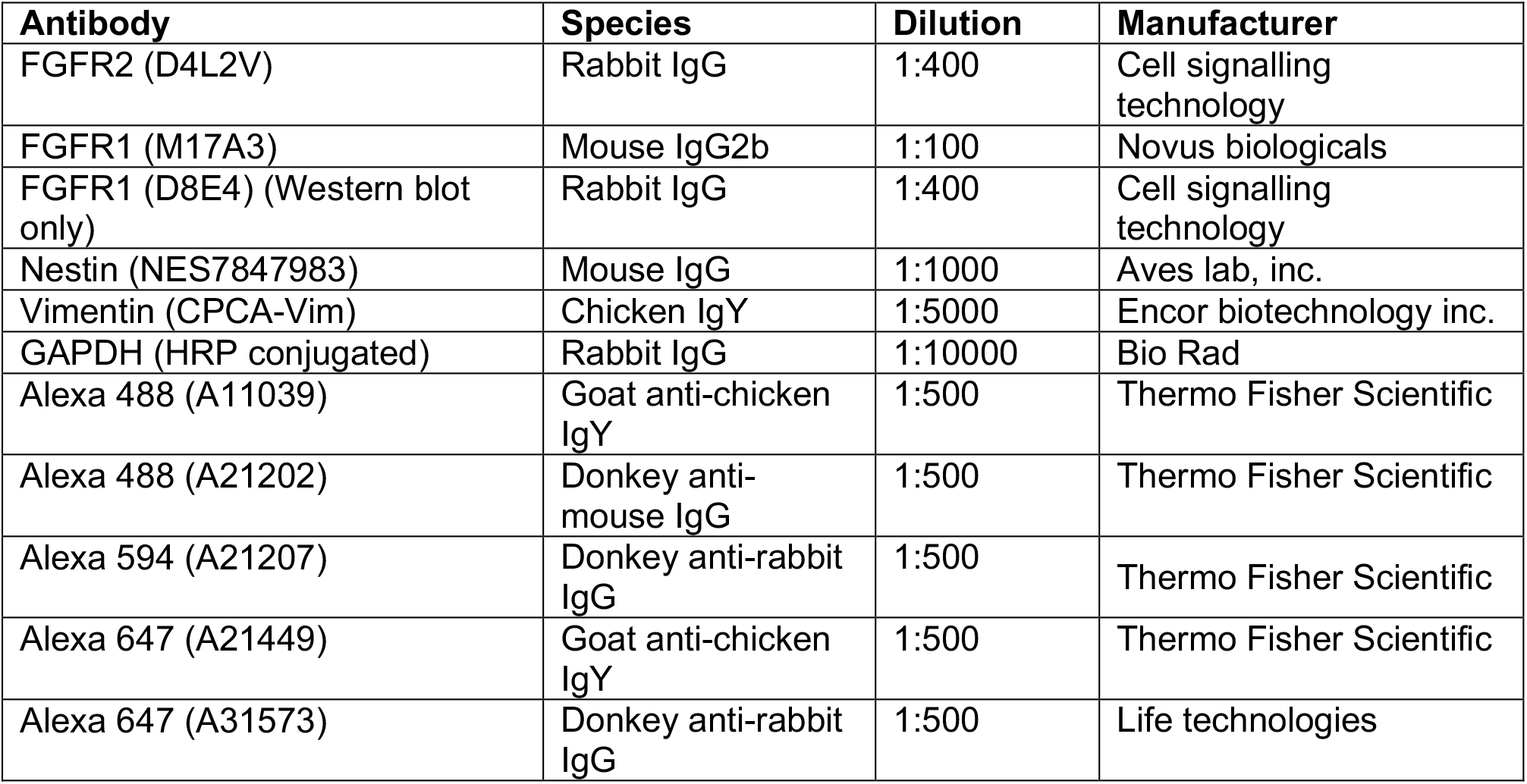
Antibodies used in this study. Table lists antibodies, clone identifier, and vendor information.

## References

1. DeAngelis LM. Brain tumors. N Engl J Med. 2001;344(2):114–123. doi:10.1056/NEJM200101113440207

2. Kao HW, Chiang SW, Chung HW, Tsai FY, Chen CY. Advanced MR imaging of gliomas: an update. Biomed Res Int. 2013;2013:970586. doi:10.1155/2013/970586

3. Lathia JD, Mack SC, Mulkearns-Hubert EE, Valentim CL, Rich JN. Cancer stem cells in glioblastoma. Genes Dev. 2015;29(12):1203–1217. doi:10.1101/gad.261982.115

4. Langen, K, Drzezga, A, Galldiks N. PET/CT in Cancer: An Interdisciplinary Approach to Individualized Imaging. Elsevier. 2018. Chapter 12; Brain Tumors. Pages 235–254.

5. Bettegowda C, Chen LC, Mehta VA. et al. Schmidek and Sweet Operative Neurosurgical Techniques: Indications, Methods, and Results: Sixth Edition. Vol. 1 Elsevier Inc., 2012. p. 669–683.

6. Gallego O. Nonsurgical treatment of recurrent glioblastoma. Curr Oncol. 2015;22(4):e273–e281. doi:10.3747/co.22.2436

7. Binabaj MM, Bahrami A, ShahidSales S, et al. The prognostic value of MGMT promoter methylation in glioblastoma: A meta-analysis of clinical trials. J Cell Physiol. 2018;233(1):378–386. doi:10.1002/jcp.25896

8. Yun Y-R, Won JE, Jeon E, et al. Fibroblast Growth Factors: Biology, Function, and Application for Tissue Regeneration. Journal of Tissue Engineering. 2010;1(1). doi:10.4061/2010/218142

9. Birzu C, French P, Caccese M, et al. Recurrent Glioblastoma: From Molecular Landscape to New Treatment Perspectives. Cancers (Basel). 2020;13(1):47. doi:10.3390/cancers13010047

10. Ozdemir-Kaynak E, Qutub AA, Yesil-Celiktas O. Advances in Glioblastoma Multiforme Treatment: New Models for Nanoparticle Therapy. Front Physiol. 2018;9:170. doi:10.3389/fphys.2018.00170

11. Mallick, S. et al. Management of glioblastoma after recurrence: A changing paradigm. Journal of the Egyptian National Cancer Institute. 2016;28(4):199–210. doi: 10.1016/j.jnci.2016.07.001.

12. Ranjan T, Howard CM, Yu A, et al. Cancer Stem Cell Chemotherapeutics Assay for Prospective Treatment of Recurrent Glioblastoma and Progressive Anaplastic Glioma: A Single-Institution Case Series. Transl Oncol. 2020;13(4):100755. doi:10.1016/j.tranon.2020.100755

13. Siebzehnrubl F. et al. The ZEB1 pathway links glioblastoma initiation, invasion and chemoresistance. EMBO Molecular Medicine. 2013;5(8):1196–1212. doi:10.1002/emmm.201302827.

14. Rahman M, Deleyrolle L, Vedam-Mai V. et al. The cancer stem cell hypothesis: failures and pitfalls. Neurosurgery. 2011;68(2):531–545. doi:10.1227/NEU.0b013e3181ff9eb5

15. Haley EM, Kim Y. The role of basic fibroblast growth factor in glioblastoma multiforme and glioblastoma stem cells and in their in vitro culture. Cancer Lett. 2014;346(1):1–5. doi:10.1016/j.canlet.2013.12.003

16. Prager BC, Bhargava S, Mahadev V. et al. Glioblastoma Stem Cells: Driving Resilience through Chaos. Trends Cancer. 2020;6(3):223–235. doi:10.1016/j.trecan.2020.01.009

17. Sutter, R., Yadirgi, G. and Marino, S. Neural stem cells, tumour stem cells and brain tumours: Dangerous relationships?. Biochimica et Biophysica Acta (BBA) - Reviews on Cancer. 2007;1776(2):125–137. doi: 10.1016/J.BBCAN.2007.07.006

18. Gupta B, Errington AC, Jimenez-Pascual A, et al. The transcription factor ZEB1 regulates stem cell self-renewal and cell fate in the adult hippocampus. Cell Rep. 2021;36(8):109588. doi:10.1016/j.celrep.2021.109588

19. Jin X, Jin X, Jung JE, et al. Cell surface Nestin is a biomarker for glioma stem cells. Biochem Biophys Res Commun. 2013;433(4):496–501. doi:10.1016/j.bbrc.2013.03.021

20. Edwards LA, Li A, Berel D, et al. ZEB1 regulates glioma stemness through LIF repression. Sci Rep. 2017;7(1):69. doi:10.1038/s41598-017-00106-x

21. Joseph JV, Conroy S, Pavlov K, et al. Hypoxia enhances migration and invasion in glioblastoma by promoting a mesenchymal shift mediated by the HIF1α-ZEB1 axis. Cancer Lett. 2015;359(1):107–116. doi:10.1016/j.canlet.2015.01.010

22. Garros-Regulez L, Garcia I, Carrasco-Garcia E, et al. Targeting SOX2 as a Therapeutic Strategy in Glioblastoma. Front Oncol. 2016;6:222. doi:10.3389/fonc.2016.00222

23. Loilome W, Joshi AD, ap Rhys CM, et al. Glioblastoma cell growth is suppressed by disruption of Fibroblast Growth Factor pathway signaling. J Neurooncol. 2009;94(3):359–366. doi:10.1007/s11060-009-9885-5

24. Jimenez-Pascual A, Siebzehnrubl FA. Fibroblast Growth Factor Receptor Functions in Glioblastoma. Cells. 2019;8(7):715. doi:10.3390/cells8070715

25. Fukai J, Yokote H, Yamanaka R. et al. EphA4 promotes cell proliferation and migration through a novel EphA4-FGFR1 signaling pathway in the human glioma U251 cell line. Mol Cancer Ther. 2008;7(9):2768–2778. doi:10.1158/1535-7163.MCT-07-2263

26. Rand V, Huang J, Stockwell T, et al. Sequence survey of receptor tyrosine kinases reveals mutations in glioblastomas. Proc Natl Acad Sci U S A. 2005;102(40):14344–14349. doi:10.1073/pnas.0507200102

27. Toedt G, Barbus S, Wolter M, et al. Molecular signatures classify astrocytic gliomas by IDH1 mutation status. Int J Cancer. 2011;128(5):1095–1103. doi:10.1002/ijc.25448

28. Ohashi R, Matsuda Y, Ishiwata T, Naito Z. Downregulation of fibroblast growth factor receptor 2 and its isoforms correlates with a high proliferation rate and poor prognosis in high-grade glioma. Oncol Rep. 2014;32(3):1163–1169. doi:10.3892/or.2014.3283

29. Yamaguchi F, Saya H, Bruner JM, Morrison RS. Differential expression of two fibroblast growth factor-receptor genes is associated with malignant progression in human astrocytomas. Proc Natl Acad Sci U S A. 1994;91(2):484–488. doi:10.1073/pnas.91.2.484

30. Kowalski-Chauvel A, Gouaze-Andersson V, Baricault L, et al. Alpha6-Integrin Regulates FGFR1 Expression through the ZEB1/YAP1 Transcription Complex in Glioblastoma Stem Cells Resulting in Enhanced Proliferation and Stemness. Cancers (Basel). 2019;11(3):406. doi:10.3390/cancers11030406

31. Gouazé-Andersson V, Delmas C, Taurand M, et al. FGFR1 Induces Glioblastoma Radioresistance through the PLCγ/Hif1α Pathway. Cancer Res. 2016;76(10):3036–3044. doi:10.1158/0008-5472.CAN-15-2058

32. Jimenez-Pascual A, Hale JS, Kordowski A, et al. ADAMDEC1 Maintains a Growth Factor Signaling Loop in Cancer Stem Cells. Cancer Discov. 2019;9(11):1574–1589. doi:10.1158/2159-8290.CD-18-1308

33. Dobin A, Davis CA, Schlesinger F, et al. STAR: ultrafast universal RNA-seq aligner. Bioinformatics. 2013;29(1):15–21. doi:10.1093/bioinformatics/bts635

34. Love MI, Huber W, Anders S. Moderated estimation of fold change and dispersion for RNA-seq data with DESeq2. Genome Biol. 2014;15(12):550. doi:10.1186/s13059-014-0550-8

35. Benjamini, Yoav, and Yosef Hochberg. Controlling the False Discovery Rate: A Practical and Powerful Approach to Multiple Testing. Journal of the Royal Statistical Society. Series B (Methodological). 1995; 57(1): 289–300. http://www.jstor.org/stable/2346101.

36. Hoang-Minh LB, Siebzehnrubl FA, Yang C, et al. Infiltrative and drug-resistant slow-cycling cells support metabolic heterogeneity in glioblastoma. EMBO J. 2018;37(23):e98772. doi:10.15252/embj.201798772

37. Krämer A, Green J, Pollard J Jr, Tugendreich S. Causal analysis approaches in Ingenuity Pathway Analysis. Bioinformatics. 2014;30(4):523–530. doi:10.1093/bioinformatics/btt703

38. Wang K, Ji W, Yu Y, et al. FGFR1-ERK1/2-SOX2 axis promotes cell proliferation, epithelial-mesenchymal transition, and metastasis in FGFR1-amplified lung cancer [published correction appears in Oncogene. 2020 Oct;39(42):6619-6620]. Oncogene. 2018;37(39):5340–5354. doi:10.1038/s41388-018-0311-3

39. Vad-Nielsen J, Gammelgaard KR, Daugaard TF, Nielsen AL. Cause-and-Effect relationship between FGFR1 expression and epithelial-mesenchymal transition in EGFR-mutated non-small cell lung cancer cells. Lung Cancer. 2019;132:132–140. doi:10.1016/j.lungcan.2019.04.023

40. Tomlinson DC, Baxter EW, Loadman PM, Hull MA, Knowles MA. FGFR1-induced epithelial to mesenchymal transition through MAPK/PLCγ/COX-2-mediated mechanisms. PLoS One. 2012;7(6):e38972. doi:10.1371/journal.pone.0038972

41. Shimizu D, Saito T, Ito S, et al. Overexpression of FGFR1 Promotes Peritoneal Dissemination Via Epithelial-to-Mesenchymal Transition in Gastric Cancer. Cancer Genomics Proteomics. 2018;15(4):313–320. doi:10.21873/cgp.20089

42. Hopkins A, Coatham ML, Berry FB. FOXC1 Regulates FGFR1 Isoform Switching to Promote Invasion Following TGFβ-Induced EMT. Mol Cancer Res. 2017;15(10):1341–1353. doi:10.1158/1541-7786.MCR-17-0185

43. Qin S, Yuan Y, Liu H, et al. Identification and characterization of sex-dependent gene expression profile in glioblastoma. Neuropathology. 2022. doi:10.1111/neup.12845

44. Huang W, Wang J, Zhang D, et al. Inhibition of KIF14 Suppresses Tumor Cell Growth and Promotes Apoptosis in Human Glioblastoma. Cell Physiol Biochem. 2015;37(5):1659–1670. doi:10.1159/000438532

45. Bittencourt LFF, Negreiros-Lima GL, Sousa LP, et al. Correction to: G3BP1 knockdown sensitizes U87 glioblastoma cell line to Bortezomib by inhibiting stress granules assembly and potentializing apoptosis. J Neurooncol. 2019;144s(3):475. doi:10.1007/s11060-019-03276-y

